# Photoperiod-Induced Neurotransmitter Switching in the Circadian Pacemaker Regulates Hypothalamic Dopamine Expression

**DOI:** 10.1101/2020.06.16.155440

**Authors:** Alessandra Porcu, Sathwik Booreddy, David K. Welsh, Davide Dulcis

## Abstract

Light, circadian clocks, and rhythmic behaviors interact closely to produce a temporal order that is essential for the survival of most living organisms. In mammals, the principal circadian pacemaker in the brain is the suprachiasmatic nucleus (SCN), which receives direct retinal input and synchronizes itself and other brain regions to the external light-dark cycle. Altered day length (photoperiod) and disrupted circadian rhythms are associated with impaired memory and mood in both humans and animal models. Prior work demonstrated that altering photoperiod can change neurotransmitter (NT) expression in the periventricular nucleus (PeVN) of the hypothalamus in adult rat brain. Here we show that neuromedin S-(NMS-) and vasoactive intestinal polypeptide-(VIP-) expressing neurons in the SCN also display photoperiod-induced neurotransmitter switching. Such photoperiod-dependent NT plasticity is retained in *Bmal1*-KO mice, indicating that NT plasticity in the SCN does not require a functional circadian clock. Utilizing a conditional viral DO-DIO vector as an historical marker of NT expression in the SCN, we further reveal that short-day photoperiod induces a cluster of *non*-NMS-expressing neurons to undergo NT switching and acquire the NMS phenotype. Selective chemogenetic activation of NMS neurons, but not VIP neurons, during the dark phase induces a significant delay in the timing of locomotor activity onset and is sufficient to increase the number of dopaminergic neurons in the PeVN. Our findings provide novel insights into molecular adaptations of the SCN neuronal network in response to altered photoperiod that affect neuronal circuit function in the hypothalamus and lead to changes in circadian behavior.

## Introduction

Seasonal changes in day length (photoperiod) affect numerous physiological functions of living organisms, particularly at high latitude^1^. Shortened day lengths in winter in northern countries are associated with seasonal affective disorder (SAD), a mood disorder characterized by depressive episodes in the fall or winter and a remission of symptoms in the spring or summer^2^. SAD patients treated with bright light therapy at dawn often display a remission of depressive symptoms within days of treatment^3^. Bright light therapy is also used to treat non-seasonal major depression, postpartum depression, and bipolar disorder^4–7^, and improves the life quality of patients with Alzheimer’s and Parkinson’s diseases^8,9^. Despite the well-known impact of light on human health, our knowledge of the mechanisms by which the brain adapts to altered environmental light is still incomplete. In mammals, intrinsically photosensitive retinal ganglion cells (ipRGCs) contain the photopigment melanopsin and are responsible for non-image-forming light detection^10,11^. A major ipRGC target is the hypothalamic suprachiasmatic nucleus (SCN), the circadian pacemaker that synchronizes to the external light-dark cycle and communicates information about it to other brain regions and peripheral tissues. Because of its central role in detecting and relaying photic information, the SCN is a key brain structure that regulates daily rhythms and seasonal physiology^12^. Previous studies showed that distinct ipRGC subpopulations drive effects of light on learning and mood^13^. Importantly, aberrant light exposure does not alter depression-like behavior or cognitive function in animals lacking ipRGCs^11,14^, indicating that light regulates these behavioral effects through ipRGCs and their target regions, including the SCN^13^. The responsiveness of the SCN to light and its ability to regulate mood and learning^15–17^ together suggest the possibility of an SCN-mediated effect of photoperiod on mental health.

The SCN is a small nucleus comprised of ∼10,000 small-diameter neurons^18,19^, nearly all of which express the neurotransmitter (NT) gamma-aminobutyric acid (GABA) in combination with various neuropeptides. The retinorecipient SCN core is made up of light-responsive neurons that receive glutamatergic input from the ipRGCs via the retino-hypothalamic tract (RHT)^20^. Vasoactive intestinal polypeptide (VIP) and/or gastrin-releasing peptide are mainly expressed in this core region^21,22^, whereas arginine vasopressin (AVP) is mainly expressed in the more dorsal shell region, which receives input from fewer RHT fibers^23^. VIP and GABA are important for synchronizing SCN neurons to one another through local synaptic connections^23,24^. In addition to the classically characterized SCN neuropeptides, neuromedin S (NMS) is a neuropeptide with SCN-restricted expression that plays an important role in SCN function. It is expressed in ∼40% of all SCN neurons, including most, but not all, SCN neurons expressing AVP and VIP^25^.

SCN neurons coordinate with one another to adapt to different photoperiods, leading to highly plastic changes at cellular and network levels^12^. Traditional forms of neural plasticity and changes in clock gene expression patterns contribute to such changes of SCN pacemaker organization^17,26–28^. For instance, photoperiod modulates the 24 hour profiles of AVP and VIP mRNA in the SCN^29^, as well as the expression of clock genes, such as *Rev-erbα, Period 2*, and *Clock*^30^. Remarkably, the SCN encodes seasonal changes in photoperiod by the relative phases of daily rhythms of electrical activity and clock gene expression among individual SCN neurons^31–33^. Seasonal changes have also been identified in human SCN. Using postmortem brain tissue obtained from young subjects, Hofman et al. found that the number of AVP- and VIP-containing neurons was significantly higher in early autumn compared to late spring/early summer^34–36^. Together these findings suggest that the SCN adapts to seasonal changes in environmental light exposure through dynamic, coordinated regulation of its neuronal network.

In addition to this photoperiod-induced neuroplasticity in the SCN network, changing photoperiod has recently been shown to alter the number of neurons expressing particular neurotransmitters in the periventricular nucleus (PeVN) of the hypothalamus. Dulcis et al. showed that seasonal changes in light exposure can induce this novel form of NT plasticity via somatostatin-dopamine switching in the adult rat^37^ and mouse^38^ hypothalamus, with concomitant changes in stress response. Summer and winter light exposure have also been associated with changes in the total number of dopaminergic neurons in humans^39^. Because changes in photoperiod induce NT plasticity in the PeVN, which is not directly retinorecipient but receives dense projections from the SCN^40^, we hypothesized that changing photoperiod may similarly alter NT expression in SCN neurons, and that this may contribute to the photoperiod-induced plasticity in the PeVN.

In this study, we exposed adult male and female mice to long-day or short-day photoperiod for 15 days. We found that mice exposed to short days showed a reduced number of VIP neurons in the SCN, but an increased number of NMS neurons. Consistent with neurotransmitter switching, the co-expression ratio of these neuropeptides was also affected. We also found that short days induced a neuronal pool^41^ of pre-existing *non*-NMS neurons to undergo NT switching and acquire the NMS phenotype. Selective chemogenetic activation of NMS neurons, but not VIP neurons, during the dark phase induced a significant delay in the timing of daily locomotor activity onset and was sufficient to increase the number of dopaminergic neurons in the PeVN.

Our findings provide a novel mechanism by which the SCN neuronal network adapts to seasonal changes in photoperiod and reveal a new role of SCN NMS neurons in regulating circadian behaviors and PeVN NT plasticity. Because NMS neurons are expressed predominantly in the SCN, our data identify promising targets for development of new treatments of neuropsychiatric disorders associated with seasonal changes in environmental light exposure.

## Results

### Long days induce reciprocal changes in numbers of VIP+ and NMS+ neurons in SCN

Because mRNA levels of several SCN NTs, including VIP, NMS, AVP, and GABA, change in response to altered light-dark cycles^42–44^, we investigated whether changes in photoperiod alter the number of neurons expressing these NTs. Adult (post-natal day, P70) GAD67-GFP female and male mice^45^, which express green fluorescent protein (GFP) under the GABAergic neuron-specific promoter GAD67, were exposed to either long photoperiod (19L:5D, 19 hours light, 5 hours dark) or short photoperiod (5L:19D) for 15 days. To control for circadian time, all mice were sacrificed at the time of activity onset as monitored by wheel-running activity. SCN neurons, identified by GABA, AVP, VIP, and NMS immunohistochemical markers, were quantified by unbiased stereology. As shown in Figure 1A-B, the number of SCN VIP-immunoreactive neurons (VIP+) was greater in long vs. short photoperiod. Quantification of a series of 6 SCN coronal sections (30 μm) collected along the anterior-posterior (A-P) axis at specific distances from Bregma revealed specific sub-regions of the SCN where this change occurred. A reciprocal change was observed for NMS+ neurons: the number of NMS+ neurons in the SCN was reduced in long photoperiod (Figure 1D-E). Furthermore, quantification of coronal sections across the A-P axis revealed that these opposite changes in VIP+ and NMS+ cell numbers occurred in the same SCN regions (−0.34 mm, -0.46 mm) (Figure 1F). This regionally specific reciprocal change in numbers of VIP+ and NMS+ neurons suggests the possibility of NT switching^41^ in a subpopulation of SCN neurons.

**Figure 1.**
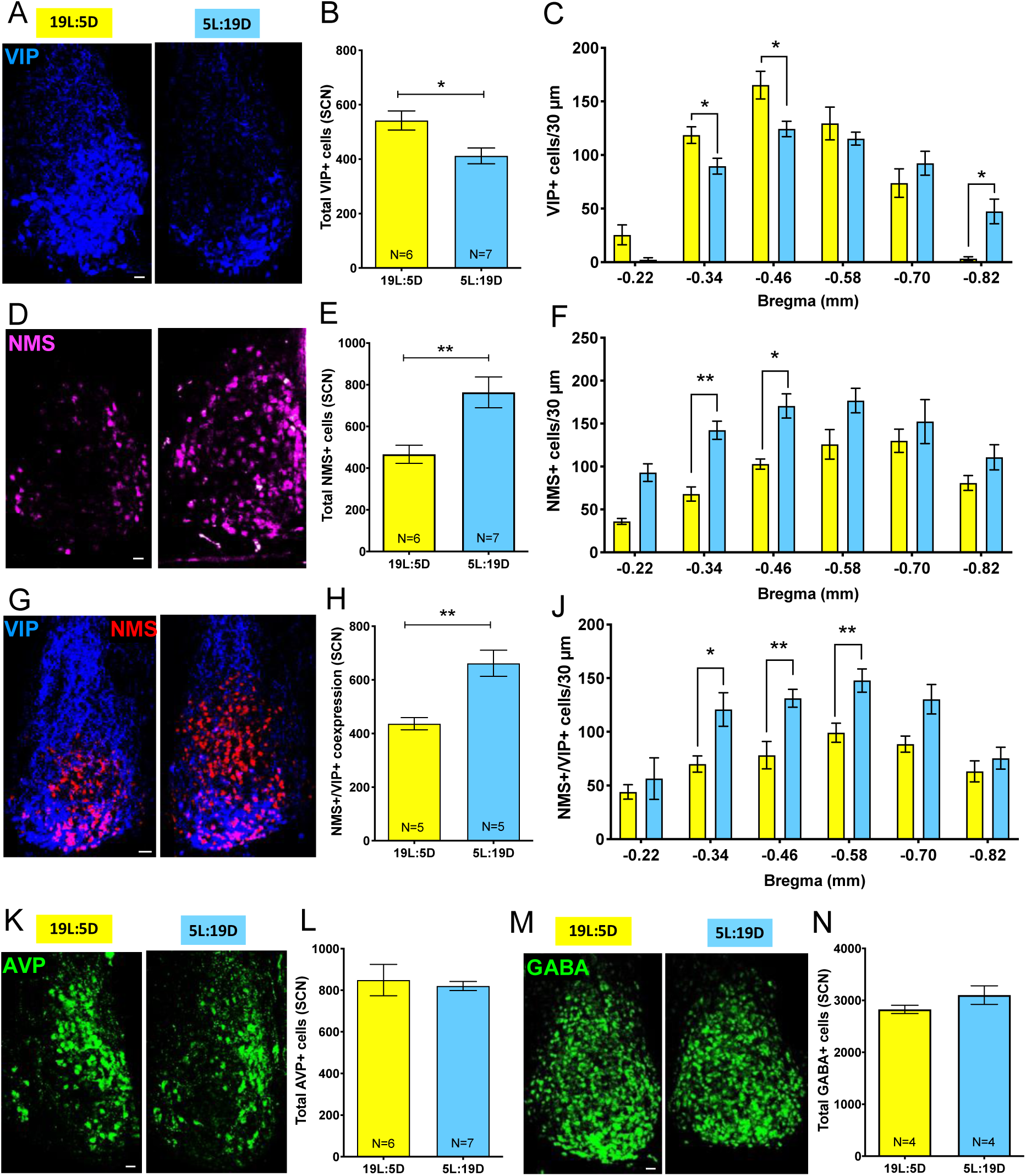
Photoperiod induces opposite effects on numbers of SCN VIP- and NMS-expressing neurons. Representative confocal micrographs of SCN from mice exposed to the long (19L:5D) or short (5L:19D) photoperiod, showing neurons identified by IHC as VIP+ (**A**), NMS+ (**D**), AVP+ (**K**), GABA (**M**), or NMS+/VIP+ co-expressing (**G**). Quantification of VIP+ (**B**), NMS+ (**E**), AVP+ (**L**), GABA+ (**N**), and NMS+/VIP+ co-expressing (**H**) neurons in the SCN following exposure to long (19L:5D, yellow bars) or short photoperiod (5L:19D, blue bars). Number of VIP+ (**C**), NMS+ (**F**), and NMS+/VIP+ co-expressing (**J**) neurons quantified per 30-µm coronal section collected along the anterior-posterior (A-P) axis at specific distances (mm) posterior from Bregma. Data are shown as means ± SEM; two-way ANOVA with Tukey’s multiple comparison post-test: *p<0.05, **p<0.01. VIP, Vasoactive intestinal polypeptide; NMS, Neuromedin S; AVP, Arginine vasopressin; GABA, Gamma aminobutyric acid. N= number of mice for each photoperiod. Scale bars: 25 µm.

To explore this possibility, we tested for co-expression of VIP and NMS in SCN neurons by double immunofluorescence. As shown in Figure 1G-H, short days significantly increased (by 34%) the level of NMS+/VIP+ coexpression, particularly in the middle region of the SCN (0.34-0.58 mm posterior to Bregma), which includes the area where we observed changes in VIP+ and NMS+ cell numbers by single immunofluorescence. This is consistent with short photoperiod inducing NMS expression in VIP+/NMS-neurons, some of which might later lose expression of VIP to become VIP-/NMS+. Finally, no differences were observed in the number of SCN AVP+ (Figure 1K-L) or GABA+ neurons across photoperiod (Figure 1M-N), indicating that NT plasticity in response to photoperiod is selective to specific cell clusters in the SCN network and does not involve the addition of new SCN neurons.

### Photoperiod-induced NT plasticity in the SCN does not require a functional circadian clock

To assess whether NT plasticity in the SCN circadian pacemaker requires clock function, we performed similar experiments in *Bmal1* knockout (*Bmal1*-KO) mice, which lack circadian rhythms in behavior^46^ or SCN neuron clock gene expression^47^. *Bmal1-KO* mice (P70) were exposed to either long or short photoperiod (19L:5D or 5L:19D) for 15 days. Mice were then sacrificed (P85), and SCN sections were processed for VIP and NMS immunohistochemistry (IHC), quantified by unbiased stereology. In *Bmal1*-KO mice, just as in GAD67-GFP mice, the number of VIP+ neurons was greater in long vs. short photoperiod (Figure 2A-B). There was also a reciprocal change for NMS+ neurons: the number of NMS+ neurons was reduced in long photoperiod (Figure 2C-D), and the decrease in NMS+ neurons was numerically similar to the increase in VIP+ neurons. These results are consistent with NT switching of neurons between VIP+/NMS- and VIP-/NMS+ phenoytpes, even in *Bmal1*-KO mice, lacking a functional circadian clock.

**Figure 2.**
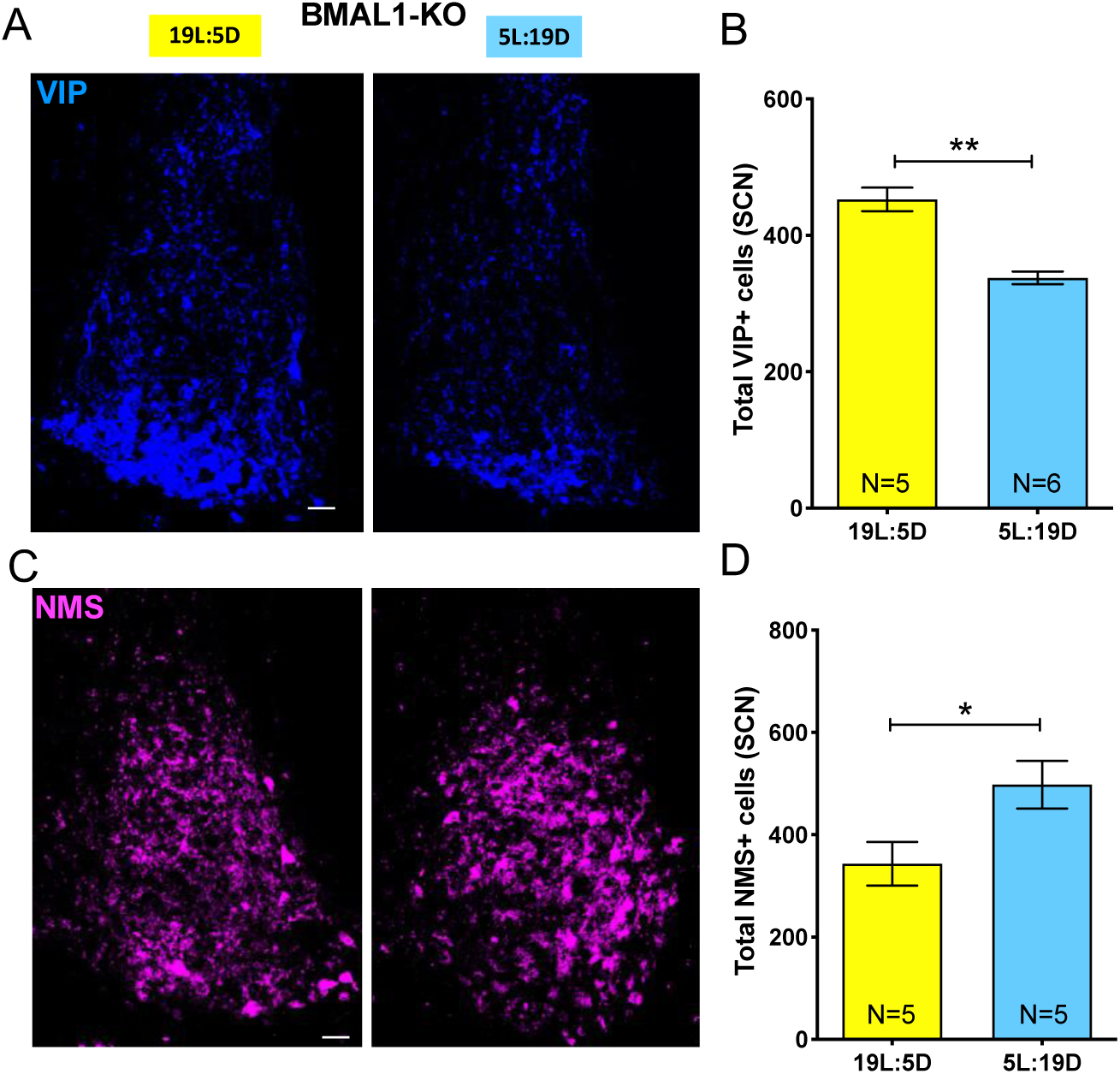
Photoperiod-induced neurotransmitter plasticity persists in *Bmal1*-KO mice. Representative confocal micrographs showing the effects of photoperiod on the number of VIP+ (**A**) and NMS+ (**C**) neurons in the SCN of *Bmal1*-KO mice. Quantification of VIP+ (**B**) and NMS+ (**D**) neurons in the SCN following exposure to long (19L:5D, yellow bars) or short photoperiod (5L:19D, blue bars). Data are shown as means ± SEM; Student *t* test *p<0.05, **p<0.01. Scale bars: 25 µm.

### Short days induce NMS expression in *non*-NMS SCN neurons

To better characterize SCN neurons exhibiting photoperiod-induced NT plasticity, we used the viral vector construct AAV-DO-DIO^48^, which acts as a Cre-dependent bistable switch, labeling cells with GFP (green, ON) in the presence of Cre recombinase, and TdTom (red, OFF) in its absence (Figure 3A). In addition, due to the stability of the GFP and TdTom reporters themselves, this construct also allows detection of cells with a history of switching between ON and OFF states, in which case cells are labeled with both reporters (yellow). By introducing AAV-DO-DIO into the SCN of NMS-Cre mice by stereotactic injection, followed by photoperiod manipulation, we aimed to distinguish NMS+ (GFP, green) and *non-*NMS (TdTom, red) neurons in SCN, as well as neurons in which switching had occurred between these two phenotypes (GFP + TdTom, yellow) in response to a change in photoperiod. Specifically, in mice subjected to short days, which in previous experiments resulted in an increased number of NMS+ neurons in SCN, we predicted an increase in number of yellow cells, reflecting new NMS expression in previously *non-*NMS (NMS-) cells.

**Figure 3.**
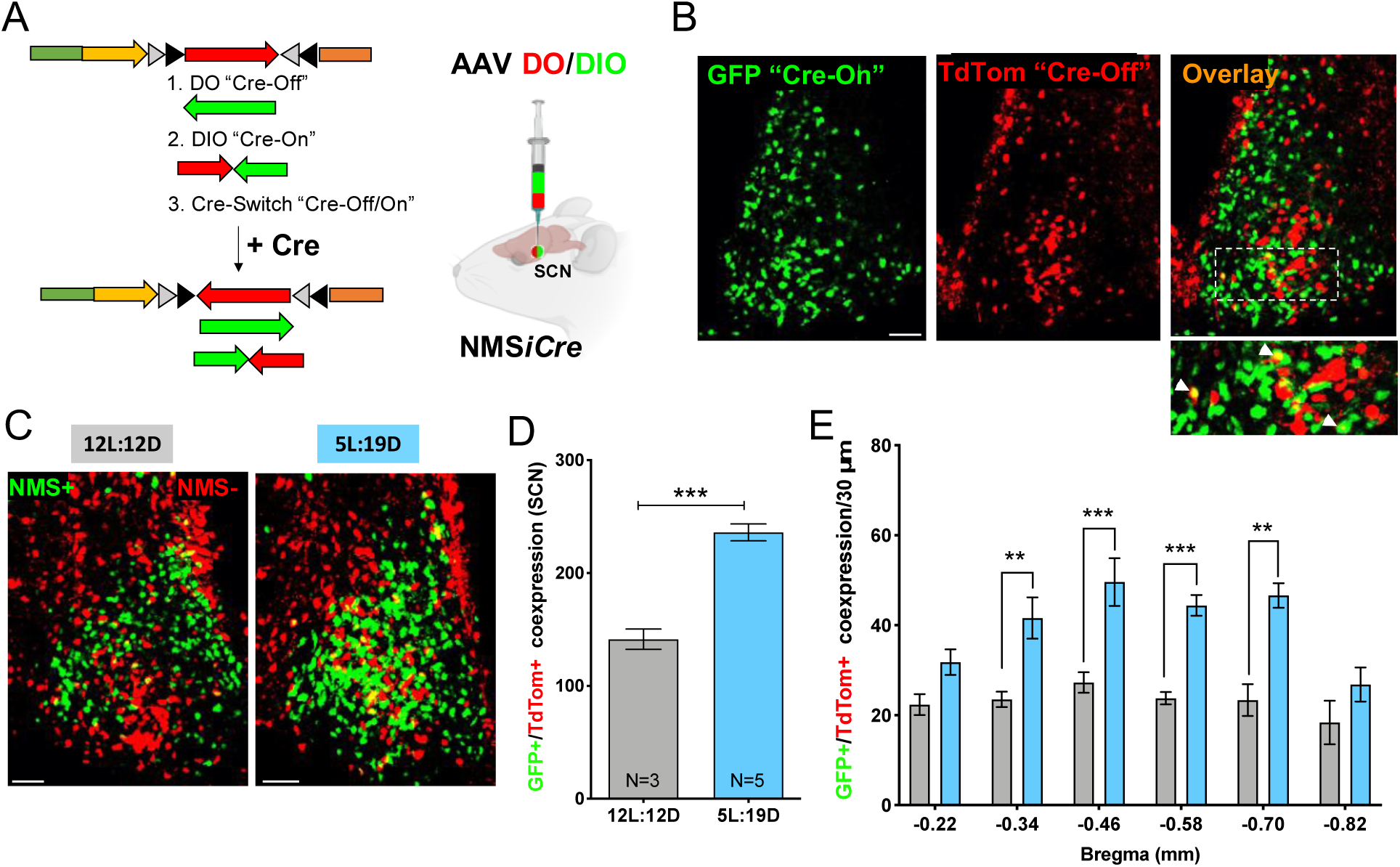
NMS plasticity in the SCN occurs by de novo expression of NMS in *non*-NMS neurons. **(A)** Genetic strategy to achieve Cre-On GFP expression and Cre-Off TdTomato expression in the SCN of NMS-iCre mice^39^. **(B)** Representative confocal micrographs showing SCN sections of an NMS-iCre control mouse (12L:12D) infected with a Cre-Switch transgene (DO-DIO) encoding Cre-On GFP (green, NMS+ cells) and Cre-Off TdTomato (red, NMS-negative cells). **(C)** Representative confocal micrographs showing the effects of short photoperiod on GFP/TdTom co-expression in DO-DIO-injected NMS-iCre mice. **(D)** Quantification of GFP/TdTom co-expression in the SCN following exposure to neutral (12L:12D, grey bar) or short (5L:19D, blue bars) photoperiod. Data are shown as means ± SEM. **(E)** Number of GFP/TdTom co-expressing neurons quantified per 30 µm coronal section collected along the anterior-posterior (A-P) axis at specific distances (mm) posterior to Bregma. Data are shown as means ± SEM; two-way ANOVA with Tukey’s multiple comparison post-test: * **p<0.01, ***p<0.001. Scale bars: 50 µm.

After stereotactic injection of AAV-DO-DIO into SCN, NMS-Cre mice raised from birth in an intermediate photoperiod (12L:12D) while allowing 2 weeks for full viral expression, were then subjected to either an additional 2 weeks of 12L:12D (control) or a short photoperiod (5L:19D). To control for circadian time, all mice were sacrificed at the time of activity onset as monitored by wheel-running activity. SCN neurons were imaged for GFP/TdTom markers and quantified by unbiased stereology. In the control condition (12L:12D), the markers revealed clear GFP (Cre-On) vs. TdTom (Cre-Off) signal separation in the SCN, reflecting distinct NMS+ (GFP, green) and NMS-(TdTom, red) phenotypes in the vast majority of SCN neurons (Figure 3B). Only a few yellow SCN neurons were detectable (GFP/TdTom co-expression, Figure 3B inset, arrowheads), reflecting a history of switching between NMS- and NMS+ phenotypes. However, in mice subjected to short photoperiod (5L:19D), which results in an increased number of NMS+ neurons in the SCN (Figure 1E), we observed a significant increase in GFP/TdTom co-expression (yellow cells, Figure 3C-D) compared to mice kept in 12L: 12D control conditions, indicating that the increase in NMS+ neurons is due to an upregulation of NMS expression in previously NMS-neurons. Furthermore, systematic quantification of GFP/TdTom co-expression along the anterior-posterior (A-P) axis of the SCN (Figure 3E) revealed that such neuronal recruitment occurred within the same middle region of the SCN where we had previously seen NT plasticity (Figure 1C, 1F, 1J). These data demonstrate that altered photoperiod can recruit pre-existing *non*-NMS neurons in the SCN to undergo NT switching and acquire the NMS phenotype.

### SCN NMS+ neurons project to the PeVN

A previous study found that dopaminergic (DA) cells in the PeVN express neuromedin receptors (NMUR2) and that these PeVN cells receive circadian information through an excitatory projection from NMS+ cells in the SCN^49^. To more fully characterize the NMS neurons in the SCN that project to the PeVN, we generated an NMS-TdTom reporter mouse line by crossing Cre-TdTom and NMS-Cre mice. Adult NMSx*iCre*-TdTomato mice (P70) were stereotactically injected with fluorescent (488 nm) retrograde beads in the PeVN (Figure 4A-B) and kept in a neutral photoperiod condition (12L:12D), then sacrificed two weeks after injection (P85). Inspection of hypothalamic coronal sections revealed that PeVN injections resulted in retrograde labeling of NMS-expressing neuron somata in the SCN (Figure 4C, arrowheads and insets 1-3). We counted ∼30 NMS+ cells per SCN section per animal, mainly in the medial SCN, that were also retrogradely labeled by beads injected into the PeVN. These data identify a subset of NMS+ neurons in the SCN which project to the PeVN, providing the connectivity that would be required for a potential regulatory role of SCN NMS neurons on PeVN DA activity in response to photoperiod-induced VIP-to-NMS switching in the SCN.

**Figure 4.**
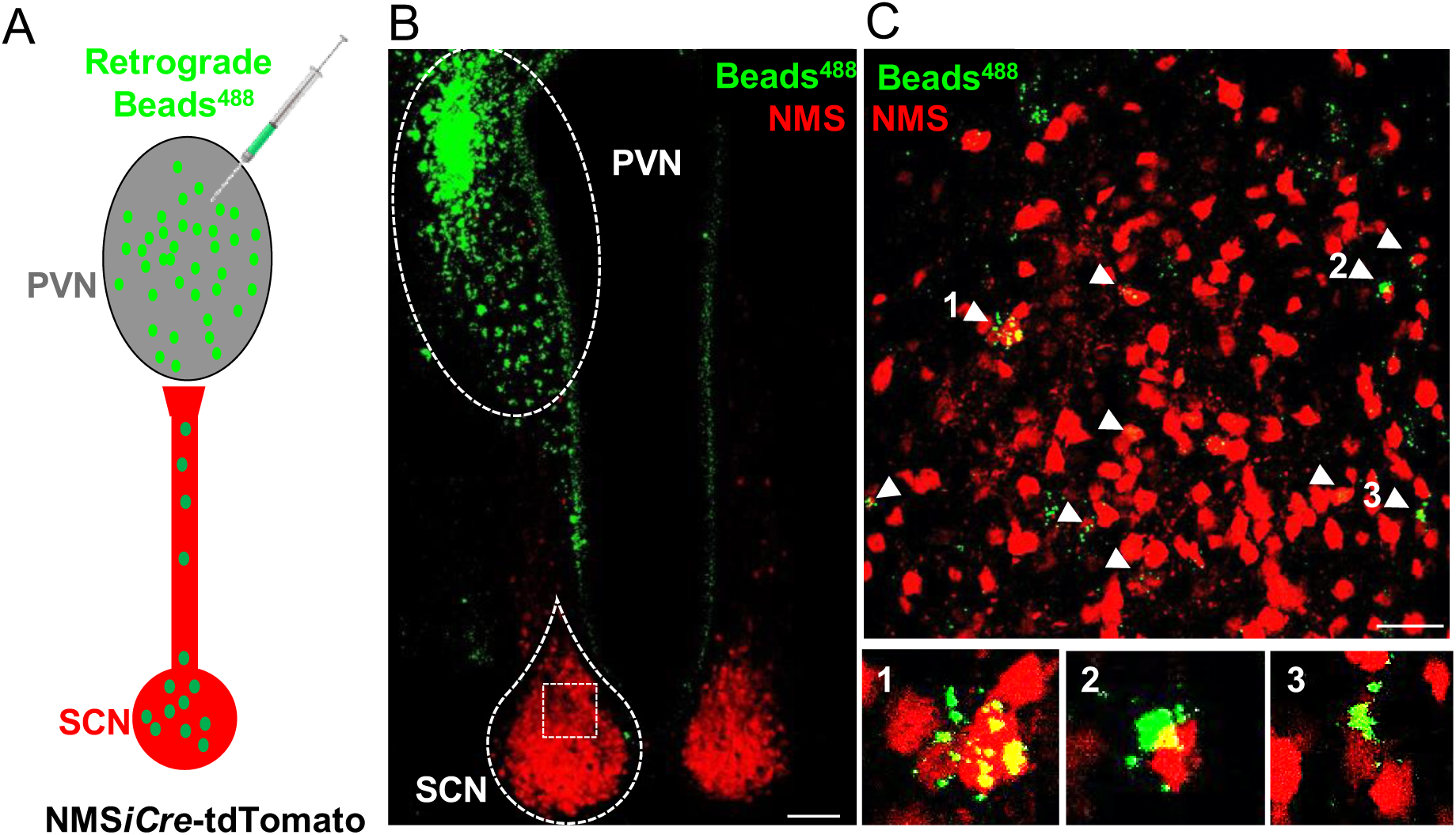
A fraction of SCN NMS+ neurons project to the PeVN. **(A)** Schematic cartoon showing the site of RetroBead^488^ injection in the PeVN and retrograde labeling of SCN somata. **(B)** Representative confocal micrograph of a hypothalamic section through the PeVN and SCN showing the RetroBead fluorescence in NMS-iCre-tdTomato mice. **(C)** Higher magnification of the SCN from the same section as in (B). Arrows indicate NMS+ (tdTomato+) and RetroBead^488^-containing somata shown at higher magnification in insets 1-3. Scale bars: 75 µm (B), 25 µm (C).

### Activation of SCN NMS+ neurons, but not SCN VIP+ neurons, delays locomotor activity onset

Earlier studies revealed that intracerebroventricular injection of NMS induced a strong increase in Fos-immunoreactive nuclei in the PeVN with a concomitant change in locomotor activity^50^. Given that PeVN DA neurons express NMS receptors and NMS depolarizes DA neurons in the PeVN^49^, we hypothesized that chronic activation of SCN NMS neurons would activate PeVN DA neurons and thereby affect circadian rhythms of locomotor activity. To test this hypothesis, we used a chemogenetic approach to chronically activate NMS+ cells in the SCN. NMS-Cre mice (P70) were stereotactically injected in the SCN with a Cre-dependent Gq-Designer Receptors Exclusively Activated by Designer Drugs (Gq-DREADDs) viral vector (Figure 5A). Two weeks after viral infusion (P75), chronic activation of Gq-DREADDs was achieved by intraperitoneal (IP) clozapine (0.01 mg/kg) injections for 15 days in a neutral photoperiod (12L:12D). Because NMS mRNA expression is higher at ZT11 (zeitgeber time 11 = 11 hrs after lights on, light phase) and lower at ZT17 (dark phase) in 12L:12D conditions^44^, IP clozapine injections were performed at ZT17, aiming to increase NMS expression during the dark phase. By prolonging daytime NMS+ neuron activation, we hypothesized that this would mimic a short-day photoperiod, thereby affecting behavioral responses. Locomotor activity was monitored using cages equipped with a running wheel and ClockLab software to generate actograms (Figure 5B). NMS-Cre mice expressing Gq-DREADDs and receiving chronic clozapine injections showed a significant delay in the time of locomotor activity onset (Figure 5C-D), whereas no effect was observed for controls: either NMS-Cre mice expressing Gq-DREADDs and receiving chronic saline injections or NMS-Cre mice expressing GFP control virus and receiving chronic clozapine injections. We then used the same Gq-DREADD approach to activate VIP neurons in the SCN of VIP-Cre mice. As shown in Figure 5E-F, these mice displayed no significant differences in timing of locomotor activity compared to control conditions. These data suggest that chronic neuronal activation of SCN NMS+ neurons but not SCN VIP+ neurons during the middle of the dark phase delays the circadian onset of locomotor activity.

**Figure 5.**
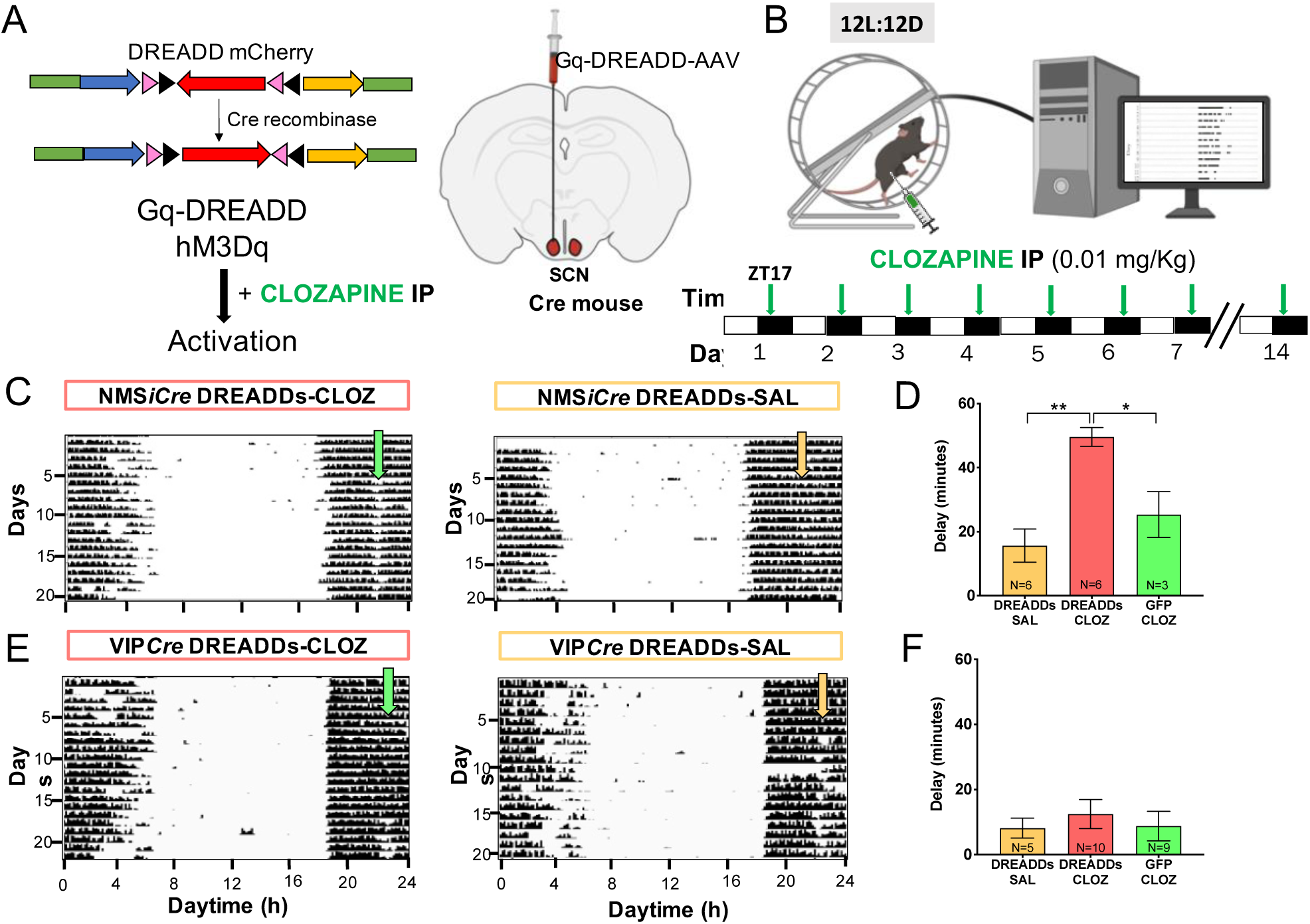
Chronic activation of NMS neurons during the dark phase delays locomotor activity onset. **(A)** Virally encoded, Cre-inducible DREADD-receptor-Cherry fusion proteins were injected into the SCN of either NMS-iCre or VIP-Cre mice. **(B)** Cartoon showing how the wheel running activity was recorded using ClockLab software to generate actograms before, during, and after daily IP clozapine (0.01 mg/kg) or vehicle (saline) injections for 2 weeks. **(C)** Representative actograms of NMS-Cre DREADD-injected mice housed in 12L:12D. The dark phase of the photoperiod is indicated with light blue shading, the light phase in white. The arrows represent the time (ZT17, zeitgeber time 17 = 17 hrs after lights on) of the first daily clozapine injection in DREADD-infected mice. **(D)** Analysis of the delay (in minutes) of wheel running activity onset was performed by measuring the difference between times of activity onset before and after clozapine injections. The first 10 onsets before clozapine injection and onsets following the first 7 injections were averaged to quantify the shift. Bar graphs show quantification of the delay of activity onset in DREADD-or control virus-infected NMS-Cre mice. Data are shown as means ± SEM; one-way ANOVA with Bonferroni’s multiple comparison post-test: *p<0.05, **p<0.01 (SAL= saline, CLOZ= clozapine, N = animals in each experimental group). (**E**) Representative actograms of VIP-Cre DREADD-injected mice housed in 12L:12D, with markings as in C. **(F)** Bar graphs show quantification of the delay of activity onset in DREADD-or control virus-infected VIP-Cre mice. Data are shown as means ± SEM; analysis and markings as in D, all comparisons non-significant.

### Chronic activation of distinct subsets of SCN neurons regulates the number of PeVN DA cells

The anatomical and functional connectivity of SCN NMS+ neurons suggest that they may have a role in regulating PeVN DA neuron activity^49^. To test whether chronic activation of SCN NMS+ neurons is sufficient to induce the kind of activity-dependent DA plasticity previously observed in the PeVN in response to short photoperiod^37,38^ NMS-Cre mice (P85) expressing either Gq-DREADDs or GFP control virus in the SCN received daily clozapine (0.01 mg/kg) or saline injections at ZT17 for 2 weeks while kept in neutral (12L:12D) photoperiod. The mice were then sacrificed (P100) for IHC analysis. SCN NMS+ neurons and PeVN DA neurons, the latter identified by tyrosine hydroxylase (TH) IHC, were quantified by unbiased stereology. As shown in Figure 6A-B, no significant differences were found in the number of SCN NMS+ neurons across experimental groups, suggesting that their Gq-DREADD-mediated activation during the dark phase was not sufficient to induce NMS plasticity in the SCN. In contrast, we found a significant increase in the number of PeVN TH+ cells in the NMS-Cre mouse group injected with Gq-DREADDs and treated with clozapine (Figure 6C-D). These results indicate that chronic activation of SCN NMS-expressing neurons during the dark phase is sufficient to recruit PeVN *non*-DA neurons to the DA phenotype.

**Figure 6.**
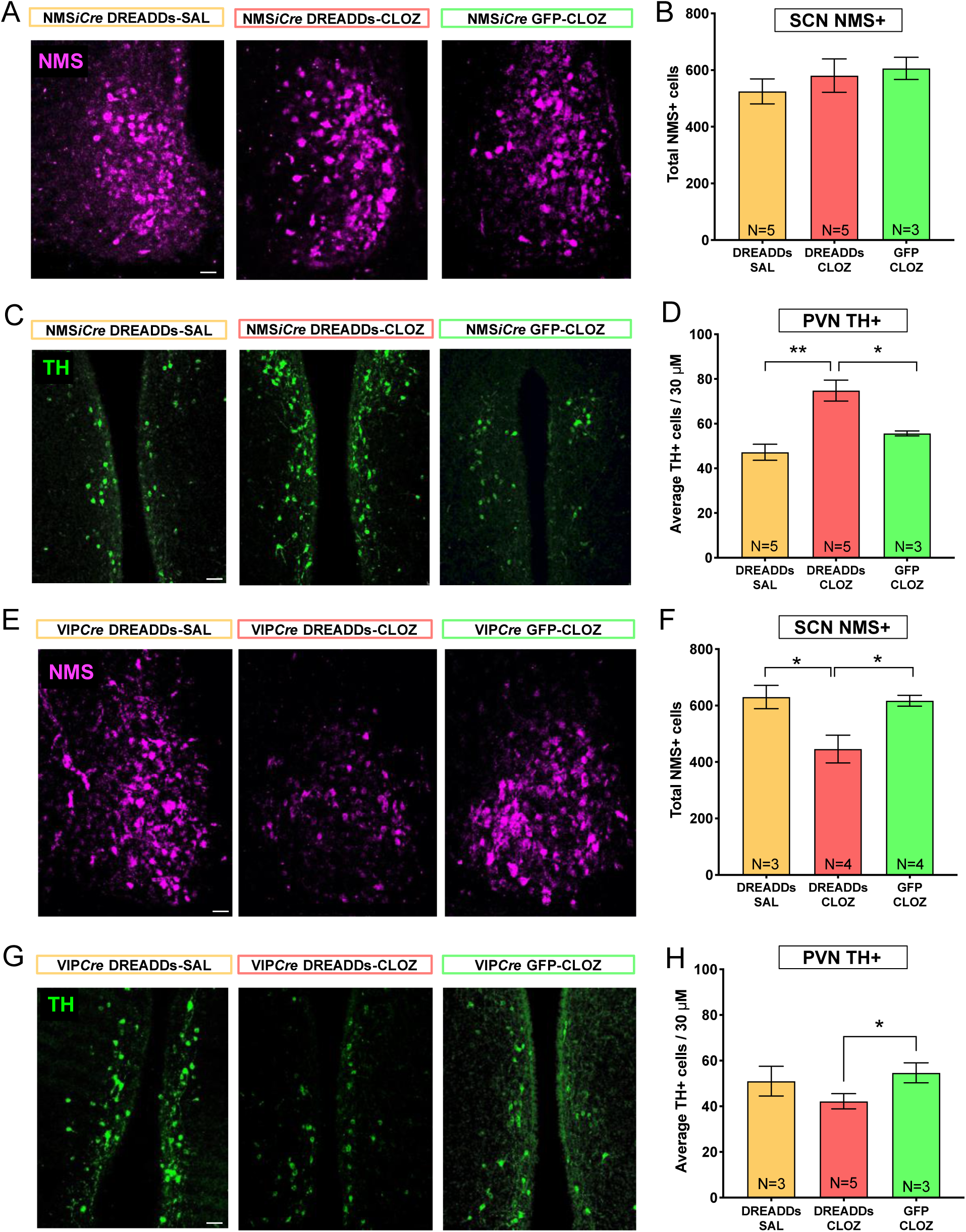
Chronic activation of SCN NMS-expressing neurons induces DA plasticity in the PeVN. **(A, C)** Representative confocal micrographs showing the effects of DREADD-mediated NMS neuron activation on the expression of NMS in the SCN (**A**) and TH in the PeVN (**C**) in NMS-iCre mice injected with either Gq-DREADD-or GFP-AAV and treated with clozapine (0.01 mg/kg) or saline. **(B, D)** Quantification of NMS+ (**B**) and TH+ (**D**) neurons for the same conditions as in A, C. Data are shown as means ± SEM; one-way ANOVA with Bonferroni’s multiple comparison post-test: *p<0.05, **p<0.01 (SAL= saline, CLOZ= clozapine, N = animals in each experimental group). **(E, G)** Representative confocal micrographs showing the effects of VIP-DREADD-induced activation on the expression of NMS in the SCN (**E**) and TH in the PeVN (**G**) in either GqDREADD-or GFP-AAV-infected VIP-Cre mice injected with clozapine or saline. **(F, H)** Quantification of NMS+ (**F**) and TH+ (**H**) neurons for the same conditions as in E, G. Data are shown as means ± SEM; one-way ANOVA with Bonferroni’s multiple comparison post-test: *p<0.05. Scale bars: 25 µm (A and E), 50 µm (C and G).

Because we found that the number of SCN VIP+ neurons also changed in response to altered photoperiod (Figure 1B), we next tested whether chronic activation of this neuronal population also affects SCN NMS or PeVN DA plasticity. VIP-Cre mice, expressing either Gq-DREADDs or GFP control virus in the SCN and undergoing a chronic treatment of daily clozapine or saline injections at ZT17 for 2 weeks, were sacrificed (P100) for IHC analysis. We found that chronic Gq-DREADD-mediated activation of SCN VIP+ neurons in the dark phase induced a significant decrease in the number of SCN NMS neurons (Figure 6E-F). In addition, we found a decrease in the number of TH+ cells in the PeVN (Figure 6G-H). These results suggest that chronic activation of VIP neuron activity during the dark phase might regulate DA plasticity in the PeVN through a concomitant decrease in number of NMS+ cells in the SCN.

## Discussion

In this study, we discovered that changes in photoperiod exposure induce a novel form of neuroplasticity in the SCN of adult mice, referred to as NT switching^41^, characterized by a NT phenotype switch between NMS+ and VIP+ neurons. Such NT plasticity in response to photoperiod was preserved in *Bmal1*-KO mice lacking a functional circadian pacemaker in the SCN. Furthermore, newly NMS-expressing neurons were recruited from a *non*-NMS pool, including a fraction of VIP+ neurons, that underwent NT switching and acquired the NMS phenotype in response to short photoperiod. Chronic activation of SCN NMS+ neurons during the dark phase delayed the daily pattern of wheel-running behavior and induced an increase in the number of PeVN DA neurons. On the other hand, chronic activation of SCN VIP+ neurons altered the number of SCN NMS+ neurons and PeVN DA+ neurons without affecting the daily wheel-running rhythms. Our findings identify a novel adaptive mechanism of the SCN network in response to altered photoperiod through which SCN neurons regulate DA function in the PeVN and circadian locomotor activity.

Previous work has shown that NT switching occurs between somatostatin (SST)- and DA-expressing neurons in the adult rat^37^ and mouse^38^ hypothalamic PeVN in response to short and long photoperiods, which mimic the seasonal changes in environmental light exposure experienced by humans at high latitudes. We discovered here that a parallel mechanism of NT switching occurs in the SCN between NMS- and VIP-expressing neurons in adult mice exposed to altered photoperiod. VIP neurons are expressed in the ventral part of the SCN and receive dense glutamatergic synaptic input from the RHT^51^, thereby playing a key role in receiving light information from the retina and relaying it to the dorsal part of the SCN. It has been shown that reorganization of the SCN circuit to adapt to changes in photoperiod requires VIP-mediated intercellular coupling^28^. *In vivo* and *in vitro* application of VIP mimics light-induced responses of the SCN^52,53^, and VIP-deficient mice do not adapt to a change in photoperiod^27^. VIP expression is influenced by environmental light, and long days induce an overall increase of VIP mRNA expression in the SCN^42^. Our findings showing an increase in the number of SCN VIP+ neurons in response to long days indicate that light can induce *de novo* VIP expression at the transcriptional and translational level in *non*-VIP neurons of the SCN. Given that the transcription factor BMAL1 is a principal driver of the molecular clock in mammals^54^, the discovery that VIP neuron recruitment was retained in *Bmal1*-deficient mice suggests that circadian clock function is not required for photoperiod-induced changes in VIP expression.

Whereas VIP expression is affected by environmental light, AVP expression is driven by the intrinsic molecular feedback loop in the core circadian clock machinery^55^. Previous studies reported changes in AVP mRNA levels in mice exposed to altered photoperiod^29^. However, we did not find any significant difference in the total number of AVP neurons in the SCN of adult mice exposed to either long or short days, indicating that light-induced NT plasticity in the SCN occurs only within specific classes of neurons such as VIP and NMS cells. Deletion of *Bmal1* alters AVP expression^56^ so profoundly that no AVP+ neurons were detectable in the SCN of *Bmal1*-KO mice (results not shown).

Another widely expressed neuropeptide in the SCN is NMS, which characterizes 40% of all SCN neurons^44^. In this study, we showed a 45% difference in the total number of SCN NMS+ neurons in mice exposed to short days, compared to mice exposed to long days. Our results are in agreement with previous studies showing changes of NMS mRNA expression induced by an altered light-dark cycle^44^. We observed reciprocal changes in the total number of VIP- and NMS-expressing neurons, indicating that light regulates VIP and NMS expression in opposite directions. This was also confirmed by the changing level of VIP/NMS co-expression across light conditions. Because only ∼20% of NMS neurons express VIP in a neutral 12L:12D photoperiod^25^, our data suggest that the increase of SCN NMS+ neurons in response to short days might occur in part by recruitment of a fraction of differentiated VIP+ neurons which downregulate their VIP expression and acquire novel NMS expression.

Using a novel viral approach (DO-DIO^48^), we uncovered for the first time temporal and spatial recruitment of new NMS-expressing neurons within the adult SCN network, according to the reserve pool hypothesis^41^. Because GFP/TdTom co-expression in the SCN significantly increased in DO-DIO-injected NMS-Cre mice exposed to short days compared to mice kept in 12L:12D, NT switching must occur by recruitment of *non*-NMS(TdTom)-expressing neurons newly acquiring the NMS phenotype. The same approach could be used to reveal whether long days induce *non*-VIP neurons to change their NT phenotype.

Given that light stimulates the release of two RHT transmitters, PACAP and glutamate^57–59^, which activate Ca^2+^signaling and CREB expression in the SCN^60,61^, the NMS+/VIP+ interchange observed in this study might represent part of a complex transcriptional response of the SCN to altered photoperiod. CREB regulates the expression of the circadian clock proteins PERIOD 1 and 2 and the neuropeptide VIP^62^, such that expression of these genes is activity-dependent^63^. Thus, light could orchestrate transcriptional events to drive NT plasticity in the SCN by altering the Ca^2+^ and CRE transcriptional pathways. The mechanism behind the light-induced NMS+/VIP+ balance in the SCN is particularly relevant given that both VIP and NMS neurons are essential components of the circadian light transduction pathway. Future studies are required to reveal the transcriptional mechanisms underlying light-induced NT plasticity in the SCN.

Lee et al. found that NMS neurons regulate SCN network function and *in vivo* circadian rhythms through synaptic transmission, suggesting that NMS might represent a key mediator of the effects of seasonal changes in day length on circadian behavior^25^. Here we showed that NMS-Cre mice expressing Gq-DREADDs selectively in the SCN NMS+ neurons and receiving IP clozapine injections at ZT17 (dark phase) showed significant delays of wheel-running activity onset. The activation of SCN NMS+ neurons at ZT17 mimicked the phase delay of wheel-running activity induced by a light pulse in the early night of nocturnal rodents^64^. No effect was observed in VIP-Cre mice expressing Gq-DREADDs. Our findings suggest an important contribution of SCN NMS+ neurons to regulation of light-dependent circadian behaviors.

SCN neurons send major output signals to the PeVN region^65–67^ via diffusible factors (including neuropeptides)^68^ and neuronal projections^69^. The PeVN is a heterogenous nucleus expressing various NTs, including DA and somatostatin, and synthesizing various hormones, such as corticotropin-releasing hormone, which regulates the stress response^70,71^. Recently it has been shown that NMS neurons, which are uniquely expressed in the hypothalamus and nowhere else in the brain, release NMS at specific PeVN target synapses to regulate the activity of DA neurons^49^ expressing NMS receptors. Using a retrograde labelling approach, we confirmed that SCN NMS neurons project to the PeVN, revealing that they possess the right connectivity to dynamically regulate PeVN DA physiology in response to seasonal changes in photoperiod. Future studies are needed to test the effect of SCN NMS+ activation on stress responses.

Previous studies found that chronic stress or altered photoperiod induces NT plasticity in the PeVN^37,72^. Specifically, populations of interneurons in the adult rat^37^ and mouse^38^ PeVN switch between DA and SST expression in response to short or long days. The readiness of neurons to display NT plasticity in response to environmental stimuli depends on their distinct Ca^2+^-mediated signaling and transcription factor signatures^73–76^, and NMS application alters Ca^2+^ signaling in PeVN DA cells^49^. The increase we observed in number of PeVN DA neurons induced by sustained DREADD-mediated activation of SCN NMS+ neurons in the dark phase reveals how chronic activation of this pathway is sufficient to induce DA plasticity in the PeVN, even without altering the number of SCN NMS+ neurons. An important clue to how light might regulate VIP expression in the SCN was offered by our DREADD-mediated activation of VIP+ neurons during the early night, resulting in a significant decrease of SCN NMS+ neurons with a parallel change in PeVN DA+ neurons, indicating that artificial activation of SCN VIP+ neurons recapitulated the light-induced changes in SCN NMS+ and PeVN DA+ neurons observed in mice exposed to long days^37,77^. Our findings begin to suggest the broad outlines of an activity-dependent mechanism through which SCN VIP+ cells are recruited to acquire an NMS phenotype that in turns affects DA plasticity in the PeVN. In summary, chronic changes in SCN neuronal activity induced by long or short days induce NT switching between NMS and VIP neurons, which in turn reorganizes the SCN-PeVN circuit (Figure 7) and produces seasonal changes in daily locomotor activity rhythms.

**Figure 7.**
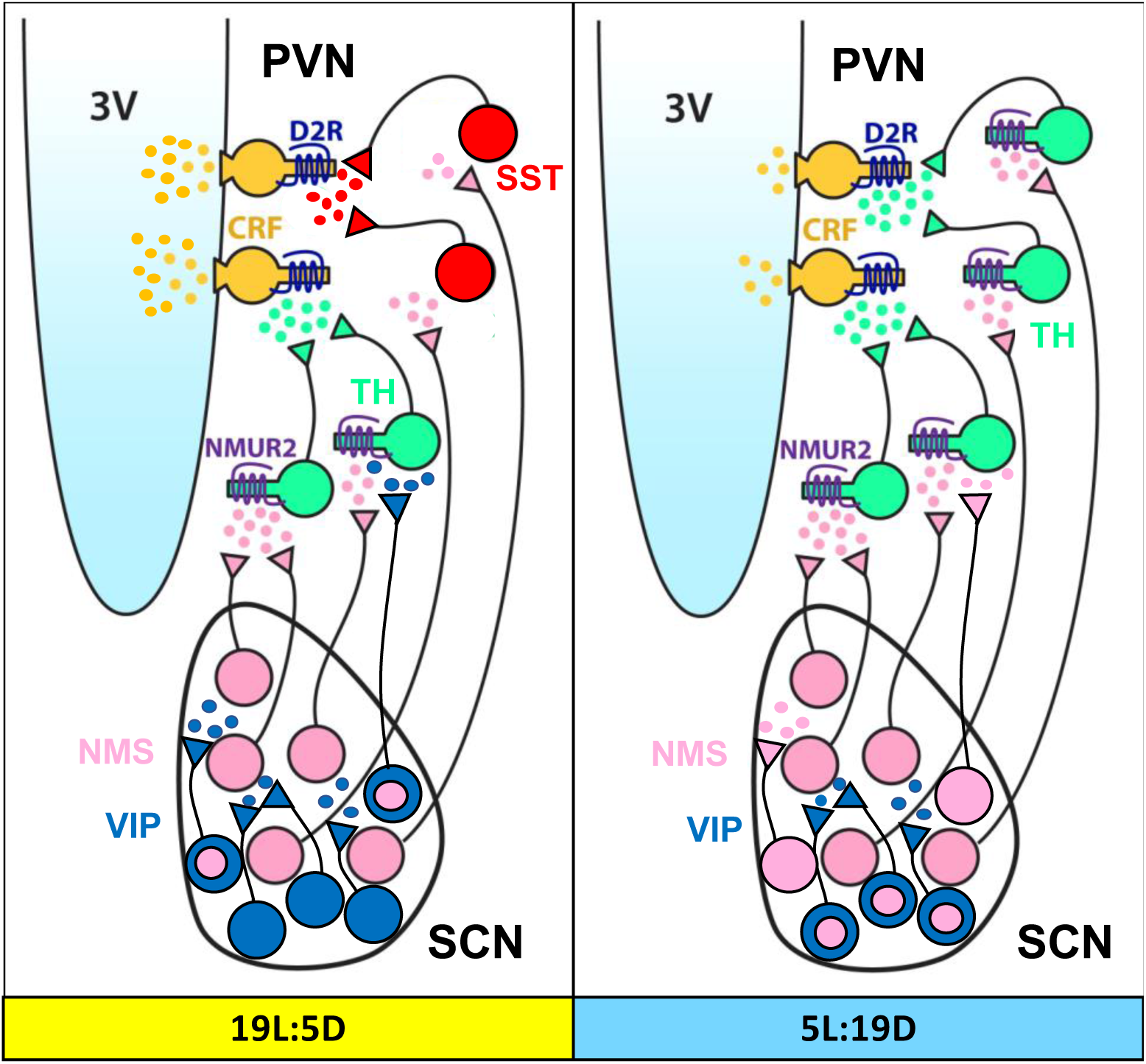
Graphic summary of photoperiod-induced NT switching in the SCN-PeVN network. In adult mice, long photoperiod (19L:5D) exposure induces an increase in the number of neurons expressing vasoactive intestinal polypeptide (VIP), whereas short-day (5L:19D) photoperiod increases the number of neuromedin S-(NMS) expressing neurons in the SCN. Co-expression of NMS and VIP neurons increases in SCN neurons of mice exposed to short days, consistent with NT switching. A parallel increase of tyrosine hydroxylase-(TH) expressing neurons in mice exposed to short days^37,77^ can also be achieved by experimental activation of NMS neurons. We propose a mechanism in which light input to the SCN regulates the number and activity of dopamine cells in the PeVN through a recruitment of new NMS neurons in the SCN from a VIP-expressing reserve pool. 3V, third ventricle; PeVN, periventricular nucleus; SST, somatostatin; SCN, suprachiasmatic nucleus; D2R, dopamine type 2 receptor; NMUR2, neuromedin S receptor; CRF, corticotropin-releasing factor. NMS/VIP co-expression is indicated by a pink circle inside a blue circle.

An important attribute of the SCN is its seasonal encoding system: a ‘memory’ for photoperiod. For example, when mice are transferred from a particular photoperiod into constant darkness (DD), distinct phase relationships among SCN neurons characteristic of that photoperiod are preserved, and mice continue to show the same characteristic behavioral patterns for days or weeks^27,32^. In addition, without VIP, the ‘memory’ of the SCN for photoperiod is lost: circadian behavior and SCN neuronal physiology appear to be unable to adapt to seasonal changes^27^. NT switching likely contributes to the mechanism through which the SCN stores seasonal light information, exerting anatomical and functional reorganization within the SCN network as well as in its target regions, including the PeVN.

This view is also supported by studies in humans showing that the volumes of the SCN and the PeVN fluctuate seasonally, with peaks in spring and autumn, and corresponding changes in the numbers of VIP and AVP neurons^34,78^. Seasonal variations in SCN morphology could also reflect the periodic incorporation of new neurons within the nucleus, although there is limited evidence of neurogenesis in the hypothalamus in adulthood. Finally, no seasonal fluctuations were found in human brain regions not involved in the temporal organization of biological processes^78^, suggesting that seasonal changes in NT expression are limited to those nuclei that are directed affected by light and transmit photoperiod-related information. The long and the short photoperiods used in this study are comparable to the ones to which humans are naturally subjected during the summer and winter months at northern latitudes. Therefore, the anatomical adaptations we observed in the SCN-PeVN network in response to changes in light exposure represents a novel model for translational research.

Shorter days during winter are associated with seasonal affective disorder and cognitive dysfunction in both humans and nocturnal animals, whereas bright light is an effective therapy for human patients with seasonal or non-seasonal depression^11,13,79–85^. Our study showing light-dependent NT plasticity in the SCN-PeVN network is a new mechanism for seasonal adaptation that might prove useful for the development of novel targets and therapeutic approaches for treating seasonal depression and cognitive impairment induced by altered light-dark cycles.

## Materials and Methods

### Mice

Mouse studies were conducted in accordance with regulations of the Institutional Animal Care and Use Committee at University of California, San Diego. Experiments involved male and female adult mice (9-16 weeks old), maintained either in 12:12 light/dark cycles (12 hours light, 12 hours dark) in long (19L:5D) or short photoperiod (5L:19D) with food and water available ad libitum. Mice were preferentially housed 4 per cage in special light-tight, ventilated chambers located in the Biomedical Sciences Building vivarium (UC San Diego). Each enclosure measures 4’ x 8’ x 4’ and contains 3 shelves each holding 16 cages. Each enclosure is constructed of laminated particle board, with 3 doors that open along the height of the enclosure. Each chamber is individually ventilated, with measured airflow of 55 air changes per hour, greatly exceeding the 10-15 minimum specified in the Guide for the Care and Use of Laboratory Animals. Mice used for locomotor activity recording were singly housed with a wheel in their home cage.

### Immunohistochemistry

Mice used for immunohistochemistry experiments were deeply anesthetized with Xylazine/Ketamine. When no response to a tail/toe pinch was present, mice were transcardially perfused with 1% phosphate-buffered saline solution first, followed by 4% paraformaldehyde solution to fix the brain tissue. The brain was then removed from the skull and kept in a 30% sucrose solution until use. Frozen brains were sectioned (30 μm) with a standard Leica Cryostat (CM1860). For each brain, 8 coronal sections encompassing the SCN and/or the PeVN were used for immunohistochemistry. Immunofluorescent labeling was performed in free-floating slices first blocked for 1 h in 0.1 M phosphate buffer (PB) containing 3% normal horse serum (NHS) and 0.2% Triton X-100. After blocking, slices were incubated with antibodies against TH (1:500, Millipore); VIP (1:1000, Immunostar); NMS 1:500, (Penninsula labs); GFP 1:500, (Invitrogen) in phosphate-buffered saline (PBS), 3% NHS, and 0.2% Triton X-100 for 24 h at 4°C with constant shaking. After three washes in PBS, slices were incubated with fluorescent-tagged secondary antibodies (AlexaFluor anti-rabbit 647 nm, anti-guinea pig 555 nm), which were all used at 1:1000 dilution. Slices were washed and mounted in gelatin on glass slides, and cover slipped with Cytoseal mounting medium (Thermo Scientific). Images were taken with a Leica confocal microscope. Stereological quantification of NMS+, VIP+ and TH+ neurons in the SCN and PeVN of the hypothalamus was performed blind with Stereologer software. The quantification data are shown as averages of the total numbers of neurons/nucleus/animal.

### Virus injection

Mice were anesthetized with 3% isoflurane and placed in a stereotactic apparatus (David Kopf Instruments, Model 900HD Motorized Small Animal Stereotaxic). Brain injections were performed during a continuous flow of 1% isoflurane. Bilateral stereotactic injections (500 nL/side) of AAV-GqDREADDs and AAV-DO-DIO particles in the SCN were achieved according to atlas coordinates (Paxinos): antero-posterior 0.46 mm; mediolateral 0.19 mm lateral; dorsoventral -5.75 mm. To allow time for diffusion of the virus, the injection needle remained immobile for 8 minutes before removal. After surgery, mice were injected with 5 mg/kg/24 h Carprofen for analgesia. For retrograde tracing, surgery was performed as described for viral injections. A volume of 200 nL of fluorescent (488 nm) RetroBeads (LumaFluor, Inc.) was unilaterally injected in the PeVN (AP: 0.85, L: 0.15, DV: -5.5) of adult mice. To allow adequate time for RetroBead transport from PeVN terminals to somata of SCN NMS neurons, mice underwent a 10 day recovery period prior to imaging.

### Data analysis

Statistical analysis was carried out with GraphPad Prism (GraphPad Software, La Jolla, CA, USA). Statistical tests for each study are indicated in the Figure legends. Data were checked for normal distribution and homogeneous variance. For normally distributed data, a parametric test was used (one-way ANOVA or Student’s t-test).

## Acknowledgments

This work was supported by the Kavli Institute for Brain and Mind (Grant No. 2012-008) to DD, by a UC San Diego Chancellor’s Research Excellence Scholarship (2018) to AP, and by a Veterans Affairs Merit Award (I01 BX001146) to DKW. We thank Anna Nilsson, I-Chi Lai, and Benedetto Romoli (Department of Psychiatry, UC San Diego) for technical support and training. All authors report no biomedical financial interests or potential conflicts of interest.

